# Quantifying nerve decussation abnormalities in the optic chiasm

**DOI:** 10.1101/633347

**Authors:** Robert J. Puzniak, Khazar Ahmadi, Jörn Kaufmann, Andre Gouws, Antony B. Morland, Franco Pestilli, Michael B. Hoffmann

## Abstract

**Objective:** The human optic chiasm comprises partially crossing optic nerve fibres. Here we used diffusion MRI (dMRI) for the in-vivo identification of the abnormally high proportion of crossing fibres found in the optic chiasm of people with albinism.

**Methods:** In 9 individuals with albinism and 8 controls high-resolution 3T dMRI data was acquired and analyzed with a set of methods for signal modeling [Diffusion Tensor (DT) and Constrained Spherical Deconvolution (CSD)], tractography, and streamline filtering (LiFE, COMMIT, and SIFT2). The number of crossing and non-crossing streamlines and their weights after filtering entered ROC-analyses to compare the discriminative power of the methods based on the area under the curve (AUC). The dMRI results were cross-validated with fMRI estimates of misrouting in a subset of 6 albinotic individuals.

**Results:** We detected significant group differences in chiasmal crossing for both unfiltered DT (p=0.014) and CSD tractograms (p=0.0009) also reflected by AUC measures (for DT and CSD: 0.61 and 0.75, respectively), underlining the discriminative power of the approach. Estimates of crossing strengths obtained with dMRI and fMRI were significantly correlated for CSD (R^2^=0.83, p=0.012). The results show that streamline filtering methods in combination with probabilistic tracking, both optimized for the data at hand, can improve the detection of crossing in the human optic chiasm.

**Conclusions:** Especially CSD-based tractography provides an efficient approach to detect structural abnormalities in the optic chiasm. The most realistic results were obtained with filtering methods with parameters optimized for the data at hand.

**Significance:** Our findings demonstrate a novel anatomy-driven approach for the individualized diagnostics of optic chiasm abnormalities.

**Highlights:**

- Diffusion MRI is capable of detecting structural abnormalities of the optic chiasm.
- Quantification of crossing strength in optic chiasm is of promise for albinism diagnostics.
- Optic chiasm is a powerful test model for neuroimaging methods resolving crossing fibers.

## 1. Introduction

The optic chiasm is a core component of the human visual system. Here the fate of the axons is decided, such that fibers from the nasal retina cross to the contralateral hemisphere, while those from the temporal retina do not cross and remain on the ipsilateral hemisphere. This partial decussation of the optic nerves guarantees that each hemisphere receives binocular input from the contralateral visual hemifield. Beyond its clinical relevance (Hoffmann and Dumoulin 2015) and its relevance as a model for neuronal pathfinding in basic neuroscience (Prieur and Rebsam 2017; Petros, Rebsam, and Mason 2008) the optic chiasm can be used as a powerful test-bed for the development of methods that allow the in-vivo-reconstruction of fiber tracts in the human brain. A common tool for this purpose is diffusion MRI (dMRI), which uses random thermal motion of water molecules (Stejskal and Tanner 1965) to identify markers of the neuronal tissue organization. This approach, initially using the Diffusion Tensor model [DT; (Basser, Mattiello, and LeBihan 1994)], was, however, proven to be confounded by tissues with a complex microstructure comprising a mixture of crossing and non-crossing nerves (Alexander, Barker, and Arridge 2002; Tuch et al. 2002). This leads to misestimations of microstructural parameters (Oouchi et al. 2007) and the underlying fiber distribution (Jones, Knösche, and Turner 2013). Moreover, this is particularly relevant in areas with crossing fibers, which may affect as much as 90% of the brain volume (Jeurissen et al. 2013), making it an important challenge for dMRI. In order to address this challenge new models were developed, such as Q-Ball imaging (Tuch 2004), Diffusion Spectrum Imaging [DSI; (Van J. Wedeen et al. 2005)] or Constrained Spherical Deconvolution [CSD; (J-Donald Tournier, Calamante, and Connelly 2007; Descoteaux et al. 2006)]. Those and other emerging approaches, however, require a sound testing model.

The optic chiasm reflects the complex microstructure of other brain structures and has the decisive advantage that (i) the ratio of crossing and non-crossing nerve fibers is a well-known ground truth and that (ii) human diseases are known, where this ratio is significantly altered. Based on previous work (Kupfer, Chumbley, and Downer 1967), we have a clear understanding that in the neurotypical case 53% of nerve fibers in the optic chiasm travel across the optic chiasm to the contralateral lateral geniculate nucleus (LGN), while 47% project to the ipsilateral LGN. The optic chiasm was investigated in several studies evaluating the accuracy of dMRI, such as the qualitative evaluation of tracking algorithms (Staempfli et al. 2007) or the quantification of fiber crossing strength in ex-vivo chiasms (Roebroeck et al. 2008). While both studies were successful in capturing qualitative features of the optic chiasm, neither provided an accurate quantitative estimation of crossing strength that was in agreement with expected values; Staempfli et al. (2007) performed qualitative analyses only and Roebroeck et al. (2008) estimated only up to 5% of nerve fibers to cross to the contralateral hemisphere.

The optic chiasm as a test-bed for differentiating crossing and non-crossing fibers can be extended further by the inclusion of known neuropathies affecting the human optic chiasm that reveal clear abnormalities. The most frequent of these rare conditions is albinism, which is associated with an enhanced crossing of the optic nerve fibers (Guillery 1986; Morland et al. 2002; E. A. H. Hagen et al. 2005). Here the line of decussation that separates the retinal ganglion cells with a crossed projection from those with an uncrossed projection and which is normally aligned with the fovea, is shifted by on average 8 degrees into the temporal retina (Hoffmann et al. 2005; E. A. H. von D. Hagen et al. 2007; E. A. H. von D. Hagen, Hoffmann, and Morland 2008). As a result, the crossing of the optic nerves is enhanced (Hoffmann and Dumoulin 2015). Recently, the first study to report group differences in chiasm tractography between albinism and controls was published (Ather et al. 2019). It used the DT model and demonstrated that dMRI can be used to identify, at the group level, differences in chiasmal connectivity between albinism and controls.

The dMRI approach can be extended beyond the DT model (Basser, Mattiello, and LeBihan 1994), by incorporating an additional Constrained Spherical Deconvolution model [CSD; (J-Donald Tournier, Calamante, and Connelly 2007)], as well as the state-of-the-art tractography evaluation: Linear Fascicle Evaluation [LiFE; (Pestilli et al. 2014; Caiafa and Pestilli 2017)], Convex Optimization Modeling for Microstructure Informed Tractography [COMMIT; (Daducci et al. 2013, 2015)], and Spherical-deconvolution Informed Filtering of Tractograms [SIFT2; (Smith et al. 2015)]. In the present study, we compared the efficacy of these methods in identifying and quantifying optic nerve fiber misrouting at the optic chiasm in albinism and its relation to fMRI-based estimates of misrouting extent.

## 2. Methods

### 2.1 Participants

Nine participants with diagnosed albinism (5 females) and eight control subjects (6 females) were recruited for the study. The controls had no neurological or ophthalmological history, normal decimal visual acuity [≥ 1.0, Freiburg Visual Acuity Test (Bach 1996)] and normal stereo vision (Lang and Lang 1988; Donzis et al. 1983). Each participant was instructed about the purpose of the study and the methods involved and gave written informed study participation and data sharing consents. The study was approved by the Ethics Committee of the Otto-von-Guericke University Magdeburg, Magdeburg, Germany.

### 2.2 Data acquisition

All MRI data were acquired with a Siemens MAGNETOM Prisma 3 Tesla scanner with *syngo* MR D13D software and a 64-channel head coil. Diffusion and functional data was acquired in separate scanning sessions. During both sessions additional T1-weighted images were acquired.

#### 2.2.1 T1-weighted data acquisition

T1-weighted images, obtained during both dMRI and fMRI scanning sessions, were collected in sagittal orientation using a 3D-MPRAGE sequence resolution [0.9 × 0.9 × 0.9 mm^3^, FoV 230 × 230 mm, TR 2600ms, TE 4.46 ms, TI 1100 ms, flip angle 7°, image matrix: 256 × 256 × 176, acquisition time 11 min 6 s; (Mugler and Brookeman 1990)].

#### 2.2.2 dMRI data acquisition

The dMRI acquisition protocol was initiated with a localizer scan, followed by a T1-weighted MPRAGE scan and two diffusion-weighted scans [one with anterior-posterior (A>>P) and the other with posterior-anterior (P>>A) phase-encoding direction]. All data was collected during a single continuous scanning session. dMRI images were acquired with Echo-Planar Imaging (EPI) [b-value 1600 s/mm², resolution 1.5 × 1.5 × 1.5 mm^3^, anterior to posterior (A>>P) phase-encoding direction, FoV 220 × 220 mm, TR 9400 ms, TE 64.0 ms, acquisition time 22 min and 24 s]. The b-value was chosen with regard to the reported range of b-values optimal for resolving two-way crossing [for single shell acquisition: 1500-2500 s/mm^2^(Sotiropoulos et al. 2013)]. Each scan was performed with 128 gradient directions, therefore the obtained diffusion-weighted data could be described as High Angular Resolution Diffusion Imaging [HARDI; (Tuch et al. 2002)] data. The high number of gradient directions, while excessive for angular contrast allowed by b-value of 1600 s/mm^2^, enhanced the effective signal-to-noise-ratio (SNR) and thus supported residual bootstrapping. This is of importance for diffusion-MRI of the optic chiasm with its reduced SNR. The gradient scheme, initially designed for 3 shell acquisition, was generated using Caruyer’s tool for q-space sampling (Caruyer et al. 2013). Due to the acquisition time constraints, however, we limited the set of directions to single shell only. Diffusion-weighted volumes were evenly intersected by 10 non-diffusion weighted volumes for the purpose of motion correction. The second diffusion-weighted series was acquired with a reversed phase-encoding direction relative to the previous scan, i.e., posterior to anterior (P>>A). Apart from that, all scan parameters were identical to those corresponding to the preceding acquisition. The acquisition of two diffusion-weighted series with opposite phase-encoding directions enhanced the correction of geometrically induced distortions (Andersson, Skare, and Ashburner 2003). Furthermore, the additional scans provided a boost of the effective SNR in the dMRI data.

#### 2.2.3 fMRI data acquisition

Functional T2*-weighted images were acquired with EPI [resolution 2.5 × 2.5 × 2.5 mm^3^; 54 axial slices covering the whole brain, FoV 210 × 210 mm, TR 1500ms, TE 30.0 ms, flip angle 70°] during the course of a different study (Ahmadi, Herbik, et al. 2019), where methods are detailed. Briefly, the fMRI scanning session consisted of 6 functional scans, 168 time-frames each, resulting in a scan length of 252 s. The dominant eye of the albinotic participants was stimulated with moving bars that appeared within a circular aperture of 9.5° radius covering either the left or the right visual hemifields (three repetitions per hemifield stimulation). All data was acquired in a single continuous scanning session.

### 2.3 Data analysis

#### 2.3.1 dMRI data analysis

##### 2.3.1.1 dMRI data preprocessing

The data was preprocessed using a combination of software toolboxes: MRtrix 3.0 (http://www.mrtrix.org/), FMRIB’s FSL 5.0.9 (https://fsl.fmrib.ox.ac.uk/fsl/fslwiki), FreeSurfer 6.0.0 (https://surfer.nmr.mgh.harvard.edu/), ANTS 2.1.0 (http://stnava.github.io/ANTs/) and MrDiffusion (https://github.com/vistalab/vistasoft/tree/master/mrDiffusion). The preprocessing of the dMRI data from DICOM involved converting it to .mif format (compatible with MRtrix) [mrconvert command], denoising of the dMRI data [dwidenoise command; (Veraart, Fieremans, and Novikov 2016; Veraart et al. 2016)], removal of Gibbs ringing in the dMRI data [dwidegibbs command; (Kellner et al. 2015)], estimation of the susceptibility induced field in the dMRI data [topup command; (Andersson, Skare, and Ashburner 2003)] using FSL (S. M. Smith et al. 2004), correction for geometry-induced, eddy current and motion distortions in the dMRI data [eddy command; (Andersson and Sotiropoulos 2016) (Andersson and Sotiropoulos 2016)], and correction for the bias field (low frequency intensity inhomogeneities caused by uneven absorption of RF power across scanned tissue) in the dMRI data [ANTS, N4 algorithm; (Tustison et al. 2010)]. Finally, the dMRI data was coregistered to the T1-weighted images aligned to the Anterior Commissure - Posterior Commissure (AC-PC) space (mrAnatAverageAcpcNifti command from https://github.com/vistasoft/vistalab). Notably, this step also required the application of an exact transformation to the gradient table, since the gradient vectors were defined in the space of the dMRI data. The T1-weighted image was segmented into white, grey and subcortical grey matter and cerebrospinal fluid using FSL [FIRST command; (Patenaude et al. 2011)]. The white matter masks were additionally manually corrected in the region of optic chiasm using the T1-weighted images (**Figure 1A**, left and middle image).

**Figure 1.**
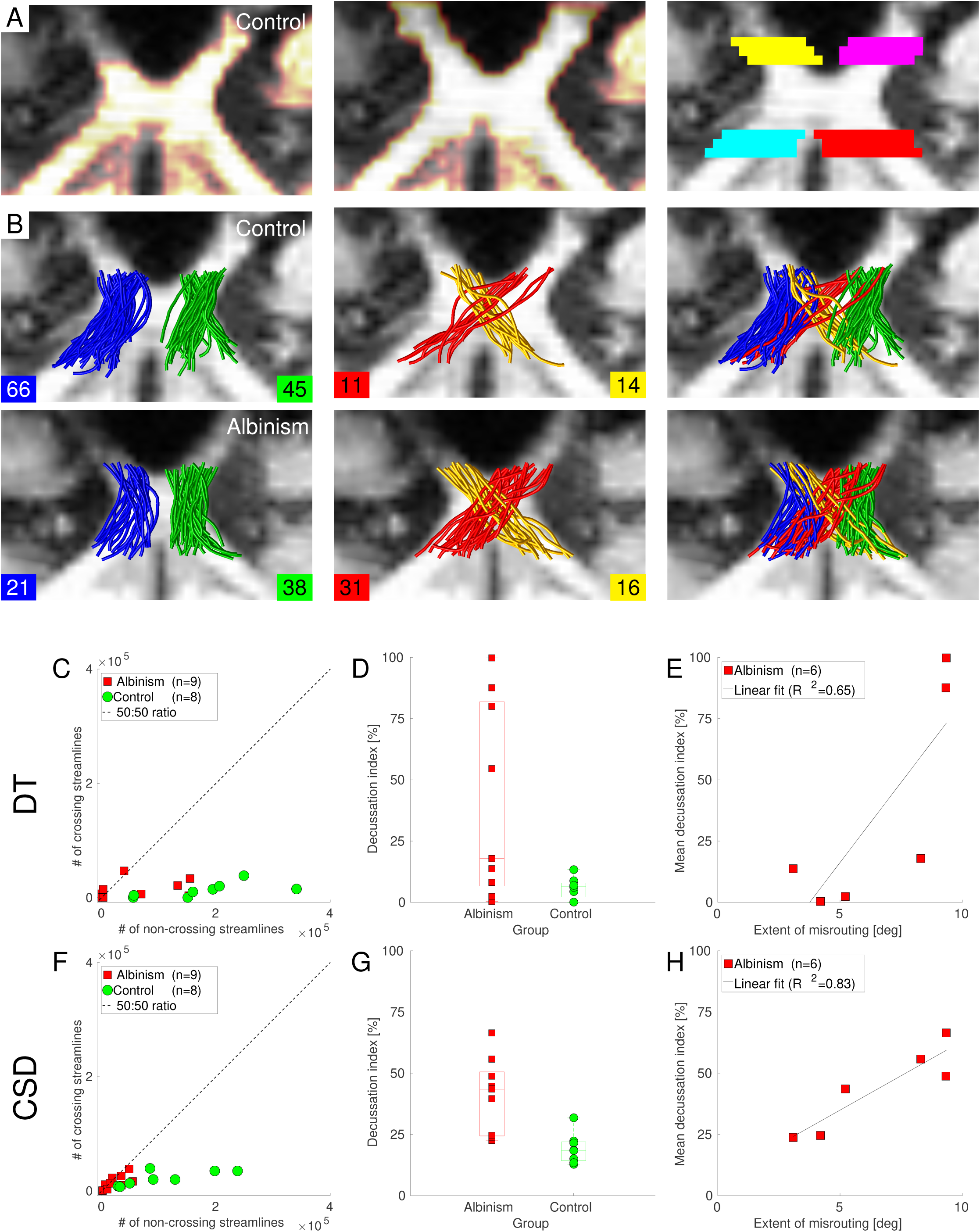
Optic chiasm – white matter mask, ROI definitions, example of tractography and quantitative results. **A.** T1-weighted image of a representative control subject overlaid with the automatically generated white matter mask (left column), manually corrected mask (middle column) and ROIs covering the intersections of the ends of optic nerves and the beginnings of the optic tracts (right column). **B.** Representative subsets of tractograms (0.125% of the total number of streamlines) created using the CSD signal model for a control (top row) and a participant with albinism (bottom row), i.e., streamlines projecting through the chiasm ipsilaterally (non-crossing; left column), contralaterally (crossing; middle column), and a combination of both streamlines (right column). **C.** Numbers of crossing vs non-crossing streamlines calculated from tractograms based on the DT model. **D.** Numbers of crossing and non-crossing streamlines calculated from tractograms based the on DT model expressed as I_D_. **E.** Correlation between the estimates of the extent of crossing obtained with dMRI using the DT model and fMRI-based pRF mapping. **F.** Numbers of crossing streamlines plotted vs non-crossing streamlines calculated from tractograms based on the CSD model. **G.** Numbers of crossing and non-crossing streamlines calculated from tractograms based on the CSD model expressed with I_D_. **H.** Correlation between the estimates of crossing obtained with dMRI, using the CSD model, and fMRI-based pRF mapping.

##### 2.3.1.2 dMRI data modeling and tractography

Two distinct diffusion signal models were applied to the dMRI data - Diffusion Tensor [DT; (Basser, Mattiello, and LeBihan 1994)] and Constrained Spherical Deconvolution [CSD; (J-Donald Tournier, Calamante, and Connelly 2007; Descoteaux et al. 2011)]. The DT model was selected in order to compare our results with previous studies that used this model alone (Roebroeck et al. 2008; Ather et al. 2019). Due to the limited performance of the DT model for populations of crossing fibers, an additional model, CSD, was also tested. The rationale behind this choice was to investigate whether the results can be improved by using a model that is more sensitive to crossing fiber populations and at the same time benefits from the high angular resolution of collected dMRI data. The modeling for DT was performed with MRtrix tool tckgen (Tensor_Prob algorithm), where dMRI data in each voxel for each streamline was residually bootstrapped, prior to DT fitting (Jones 2008), such that tracking along the principal eigenvectors of tensors was probabilistic. For the purpose of CSD modeling, an estimation of a response from voxels containing only single fiber population (single fiber response, SFR) was acquired using Tournier’s approach [dwi2response tool with -tournier option (J-Donald Tournier, Calamante, and Connelly 2013)] for a maximum harmonic order (L_max_) of 6. The fibre orientation distribution functions [fODFs; (Jeurissen et al. 2014)] were obtained for 3 different maximum harmonic orders L_max_ = 6, 8 and 10 (command dwi2fod using msmt_csd algorithm, as its hard constraints prevents the estimation of negative peaks).

For the purpose of tracking for both DT and CSD, four regions of interest (ROIs) were manually drawn on T1 images for each individual data set - two covering intersections of left and right optic nerves and two covering intersections of left and right optic tracts. ROIs were placed to be as close to the optic chiasm as possible without intersecting it (**Figure 1A**, right image). ROI widths and heights (in coronal view) had the minimal values required to fully cover the optic nerves or tracts, and each of the ROIs was 3 voxels (4.5 mm) thick to ensure proper termination of streamlines during tractography. In total there were four pairs of ROIs connecting optic nerves and optic tracts. For each model, tracking was performed between 2 ROIs, one of two optic-nerve ROIs and one of the two optic tract ROIs (**Figure 1B**), such that the created streamline groups could be either classified as crossing or non-crossing. For each pair of ROIs the streamlines were generated twice, with a switched definition of seed and target ROI, and merged into a single group of streamlines, such that the tracking results were not biased towards the direction of tracking. The tracking was limited to previously obtained and manually corrected white matter masks [following Anatomically-Constrained Tractography framework; (Smith et al. 2012)]. In addition, to reduce a potential bias in tractography caused by the specific choice of tracking parameters, Ensemble Tractography (Takemura et al. 2016) was used for all analyses. Accordingly, the tractography was performed multiple times, with each run using a different set of parameters. The modified parameters were, specifically: fractional anisotropy threshold (FA_thresh_) and maximum curvature angle between consecutive tracking steps (Curv_max_). In case of CSD this list was extended by the maximal harmonics order (L_max_) of the fitted fODFs. Notably, the CSD used the SFR obtained for only a single L_max_=6. The values of the parameters applied are summarized in **Table 1**.

**Table 1.** Tractography parameters. Sets of parameters used in generating DT- (left column) and CSD-based (right column) tractograms. The rows correspond to different parameters, from top to bottom, respectively, the cut-off FA threshold (FA_thresh_), maximum angle between consecutive tracking steps (Curv_max_), maximal harmonic order of SFR (SFR L_max_) and fODF (fODFs L_max_).

|  | DT | CSD |
| --- | --- | --- |
| $FA_{\text{thresh}}$ | 0.04,<br>0.08 | 0.04,<br>0.08 |
| $Curv_{\text{max}} [^\circ]$ | 30, 45,<br>60 | 30, 45,<br>60 |
| SFRs $L_{\text{max}}$ | - | 6 |
| fODFs $L_{\text{max}}$ | - | 6,8,10 |

For each subject, for a given combination of parameters the tractography was performed for the 4 distinct seed-target ROI pairs, and for each pair of ROIs it was performed twice (reversal of seed-target definitions), with 139 000 seeds (random locations within the seed ROI, which served as starting points for tracking) and 1000 tracking attempts per seed. This resulted in a total of 6.672*10^9^ tracking attempts per subject for the DT-based analysis and 20.016*10^9^ attempts for the CSD-based analysis. The DT-based tracking used the Tensor_Prob algorithm (Jones 2008), the CSD-based was performed with the iFOD2 algorithm (J-D Tournier, Calamante, and Connelly 2010).

The tracking with a fixed number of attempts resulted in unevenly populated streamline groups, where the count depended on the overall success ratio of tracking for (Smith et al. 2012)given set of parameters and ROIs. The tractograms were therefore biased toward the underlying structural connectivity, which allowed them to be used for the estimation of crossing strength. Furthermore, the resulting tractograms can be used as an input to filtering, where the input streamlines are modeled forward in order to explain measured diffusion signal. This, in turn, allows for the estimation of the contribution of each generated streamline to the original signal, thus addressing the stochasticity of tracking. It is of interest to investigate how well filtering performs, if the initial tractogram does not accurately represent the underlying microstructure, i.e., when it contains high-levels of noise. In order to answer this question, a second set of tractograms was created, where all the parameters were identical with the previous tractography, except for the restriction to a fixed number of streamlines per combination of groups and parameters. Precisely, the generated tractograms, further referred to as “streamline number targeted” (SNT) tractograms, were restricted to the generation of 139 streamlines or reaching 139 000 seeds with 1000 attempts per seed, with the latter condition preventing excessively long computations in cases of very low success ratio of tractography.

##### 2.3.1.3 Tractography filtering

To further investigate the robustness of the results on intra-study level, tractograms, both normally generated and SNT tractograms, were filtered with 4 separate algorithms: (1) Linear Fascicle Evaluation [LiFE; (Pestilli et al. 2014; Caiafa and Pestilli 2017; Takemura et al. 2016)], (2) Convex Optimization Modeling for Microstructure Informed Tractography [COMMIT-SZB; (Daducci et al. 2013, 2015)] using the Stick-Zeppelin-Ball model, (3) COMMIT using the Stick-Ball model [COMMIT-SB; (Daducci et al. 2015, 2013)], and (4) Spherical-deconvolution Informed Filtering of Tractograms [SIFT2; (Smith et al. 2015)]:

**(1) LiFE** (Pestilli et al. 2014; Caiafa and Pestilli 2017; Takemura et al. 2016; Avesani et al. 2019). LiFE evaluates the individual streamline paths by scoring their contribution (expressed as weights assigned to each individual streamline) to predict the measured diffusion signal across the brain white matter voxels; good streamlines positively contribute to predicting the dMRI signal (non-zero weights, higher value represents higher contribution), poor streamline paths do not contribute positively to predicting the measured dMRI signal (zero weights).
**(2) COMMIT-SZB** (Daducci et al. 2013, 2015) using a Stick-Zeppelin-Ball model. The COMMIT framework follows a similar rationale as LiFE (as well outputting weights for the whole tractogram), extending the range of model parameters that can be used for predicting the dMRI signal (i.e., it adds additional parameters in order to model both intra- and extra-axonal cellular contributions to the dMRI signal prediction). The Stick-Zeppelin-Ball model, as in (Panagiotaki et al. 2012), describes one anisotropic intra-axonal (here implemented with a tensor with axial diffusivity equal to 1.7×10^-3^ mm^2^/s and null radial diffusivity), one anisotropic extra-axonal (implemented with axial diffusivity equal to 1.7×10^-3^ mm^2^/s and radial diffusivity calculated as such, that according to tortuosity model the intra-cellular volume fractions are equal to 0.7) and one isotropic extra-axonal (implemented with tensors with isotropic diffusivities equal to 1.7×10^-3^ and 3.0×10^-3^ mm^2^/s) compartment contributing to modeling and predicting the dMRI signal for each segment of analyzed streamlines. This model was chosen due to its overall good previous performance, as demonstrated in Panagiotaki et al (2012).
**(3) COMMIT-SB** (Daducci et al. 2013, 2015) using a Stick-Ball model. Although simpler, this multi-compartment model is a more fitting match for our acquired single shell dMRI data. While the single shell data allows for the discrimination between anisotropic and isotropic contributions to the signal, it severely limits the distinction of extra- and intra-axonal signals. Therefore a model using one anisotropic intra-axonal compartment and one isotropic extra-axonal compartment is expected to be optimal for our data and was implemented using the corresponding components from the COMMIT-SZB approach.
**(4) SIFT2** (Smith et al. 2015). SIFT2 filters a tractography solution by assigning weights to the streamlines in order to optimize a model that matches the densities generated from the weighted streamline counts with the size of the modelled fiber Orientation Distribution Functions (fODFs) estimated from the signal within the individual voxels (Smith et al. 2015). While the filtering used only a subset of whole-brain tractograms and a subset of voxels from diffusion-weighted images, the information was complete, i.e., tractograms covered all anatomically connecting regions and all white matter voxels within the region we analyzed. This feature of input tractograms allows for the application of filtering – if this criterion is not fulfilled, filtering leads to erroneous results, i.e. is not applicable. It should be noted that two of the methods, i.e. LiFE and SIFT2, were applied with their default parameters, while in the case of COMMIT a wider range of parameters was tested. This, in combination with the established knowledge, demonstrates that the better the model fit to the data, the better the evaluation (Pestilli et al. 2014; Takemura et al. 2016; Rokem et al. 2015, 2017).

#### 2.3.2 fMRI data analysis

The fMRI data analysis is detailed in (Ahmadi, Herbik, et al. 2019). Briefly, the functional data was preprocessed using FreeSurfer 6.0.0, FMRIB’s FSL 5.0.9, Vistasoft (https://github.com/vistalab/vistasoft) and a toolbox developed by Kendrick Kay (https://github.com/kendrickkay/alignvolumedata). The T1-weighted images were automatically segmented into white matter volume and cortical surface using FreeSurfer. The fMRI data was corrected for motion with FSL, averaged across runs for each participant and subsequently aligned to the T1-weighted image with Vistasoft and Kendrick Kay’s alignment toolbox. The cortical surface reconstruction at the white and grey matter boundary as well as rendering of the smoothed 3D mesh (Wandell, Chial, and Backus 2000) was performed with Vistasoft.

The estimation of the pRF properties, the delineation of the visual areas, and the visualization on the smoothed mesh surface were performed using Vistasoft, as described in (Ahmadi, Fracasso, et al. 2019; Ahmadi, Herbik, et al. 2019). Polar angle and eccentricity maps were extracted and the misrouting extent was measured by calculating the mean eccentricity value [in degrees of visual angle] for the most eccentric abnormal representation in a ROI drawn at the fundus of the calcarine sulcus that coincided with the representation of the horizontal meridian. The maximal extent that could be determined was limited to the stimulus size of 9.5°. In accordance with the well-known variability of misrouting in albinism, particularly evident in cortical measures of the representation in albinism (Hoffmann et al. 2003, 2005), these values ranged between 3.0° - 9.3° (Ahmadi, Herbik, et al. 2019).

### 2.4 Quantitative and statistical analysis

The crossing strength was expressed using either the weights provided by tractogram filtering or, in the case of unfiltered tractograms, the number of the obtained streamlines (which is equivalent to the assumption that all weights are equal). For both metrics, the crossing strength was described by a pair of values - sums of streamlines/weights of streamlines that cross at the optic chiasm to the other brain hemisphere (crossing streamlines) or that remain on the same side (non-crossing streamlines). In order to reduce the dimensionality of the results, as well as to allow for cross-study comparison (Ather et al. 2019), the decussation index (I_D_) was calculated:

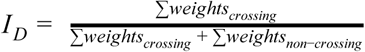

where the weights_crossing_ and weights_non-crossing_ for each streamline were either used according to the filtering results (for filtered tractograms) or set to 1 (for unfiltered tractograms).

The I_D_s obtained for each group (control and albinism), signal model (DT or CSD) and applied filtering model (none, LiFE, COMMIT-SB, COMMIT-SZB, and SIFT2) were tested for normality using the Kolmogorov-Smirnov test. As all the data samples were normally distributed, the equality of means was tested with one-tailed, two-sample t-tests at an alpha level of 5%. Additionally, for better comparability of our estimates, we calculated mean, median and standard deviation values of I_D_s.

The calculated I_D_s were subsequently entered into a Receiver Operating Characteristic (ROC) curve analysis, which returned the Area Under Curve (AUC) measure for a given combination of signal model (DT or CSD) and applied various filtering methods (none, LiFE, COMMIT-SB, COMMIT-SZB, and SIFT2). It should be noted that the estimated values were derived from the classification of the training data. As such, the obtained AUCs reflect the ability to separate already measured groups.

Finally, we tested whether the variability in the fMRI-based extent of misrouting is related to the estimates of misrouting derived from dMRI. This was tested by correlating the I_D_s derived from dMRI with the fMRI-based extent of misrouting. The results were expressed with the coefficient of determination R^2^, which describes the proportion of variance in the dependent variable (dMRI-based I_D_) explained by the independent (fMRI-based extent of misrouting). It should be noted that the fMRI-based eccentricity measures were available only for a subset of six albinotic participants.

## 3. Results

In order to compare different dMRI-based approaches to assess optic chiasm integrity, we assessed (i) non-validated DT-based tractograms, (ii) non-validated CSD-based tractograms, (iii) the impact of filtering techniques on the identification of chiasmal abnormalities at both the group and the individual level and (iv) its robustness to noise in the tractograms.

### (i) Efficacy of non-validated DT-based tractograms for the detection of chiasma abnormalities

In analogy to Ather et al. (Ather et al. 2019) we quantified the crossing strength by comparing the streamline counts for crossing and non-crossing streamlines as depicted in **Figure 1B** and **C**. In order to compare different dMRI-based approaches to assess optic chiasm integrity, we further expressed the extent of crossing at the optic chiasm in the albinotic and the control group via I_D_ (**Figure 1D**) as detailed in Methods. In accordance with Ather et al. (2018), our results demonstrated a significant I_D_ difference between the albinism and the control group [t= −2.43; p=0.014; median (mean±SEM) I_D_: 17.9% (40.5%±40.1) and 6.4% (5.8%±4.4), respectively]. A ROC analysis to assess the accuracy of the detection of chiasmatic abnormalities at an individual level, yielded an AUC of 0.61, i.e., short of that reported by Ather et al. (AUC=0.73).

### (ii) Comparison of non-validated CSD- and DT-based tractograms

The CSD-based approach also allowed for the detection of differences in the chiasm crossing at the group level [t = −3.78; p=0.0009; median (mean±SEM) decussation index 43.5% (41.1%±15.2) and 18.5% (19.2%±6.2)], between the albinism and the control group, respectively as depicted in (**Figure 1 F** and **G**). Importantly, the ROC analysis yielded a higher AUC, i.e. 0.75, than the DT-based approach, which underlines a greater discriminative power at the individual level for CSD derived streamlines. Interestingly, I_D_s vary across the albinotic subjects, for both the CSD and for the DT model. We now tested whether this variability in I_D_ across albinotic subjects was related to the well-known variability of the extent of misrouting in albinism, which is particularly evident in the cortical measures of the representation abnormality in albinism (Hoffmann et al. 2003, 2005). For this purpose, we correlated the dMRI-derived I_D_ of six albinotic participants with the strength of their misrouting as determined from fMRI-based pRF mapping [see Methods]. There was a clearly significant positive correlation for the CSD model (R^2^ =0.83; p=0.012; **Figure 1H**), while there was only a non-significant trend of a positive correlation for the DT model (R^2^=0.65, p=0.052; **Figure 1E**). This indicates a remarkable precision of the dMRI-based detection of misrouting in reflecting the extent of misrouting in albinism.

### (iii) Relevance of tractography filtering

While the previous analyses were exclusively performed for unfiltered tractograms, we now assessed the effect of tractography filtering as detailed in Methods for the (a) DT and the (b) CSD modeling approaches. As for the above analyses, their outcome is quantified via the ROC-derived AUC values and the R^2^ values for the correlation of ID and fMRI-based extents of misrouting (**Table 2** and **Supplementary Figure A.1**).

**Table 2.** Results derived from original and filtered tractograms. The rows describe different filtering techniques. The columns describe separate estimates including the p-values derived from statistical testing of equality of the I_D_s for albinism and controls. Each column is divided into two sub-columns, corresponding to the two signal models, i.e., DT and CSD.

| | mean $I_D$ for albinism [%] | | mean $I_D$ for controls [%] | | p-value for $I_D$ | | AUC | | $R^2$ | |
| --- | --- | --- | --- | --- | --- | --- | --- | --- | --- | --- |
|  | DT | CSD | DT | CSD | DT | CSD | DT | CSD | DT | CSD |
| **Unfiltered** | 40.5 ± 40.1 | 41.1 ± 15.2 | 5.8 ± 4.4 | 19.2 ± 6.2 | 0.014 | 0.00009 | 0.61 | 0.75 | 0.65 | **0.83** |
| **LIFE** | 37.0 ± 33.4 | 42.3 ± 10.0 | 11.0 ± 7.1 | 35.9 ± 5.4 | 0.024 | 0.0635 | 0.60 | 0.74 | 0.52 | 0.51 |
| **COMMIT - SB** | 42.0 ± 27.2 | 41.3 ± 6.0 | 20.3 ± 8.4 | 28.6 ± 4.2 | 0.024 | 0.00008 | 0.56 | 0.94 | 0.34 | 0.32 |
| **COMMIT- SZB** | 46.3 ± 29.8 | 44.8 ± 8.4 | 22.5 ± 11.0 | 30.6 ± 5.0 | 0.025 | 0.0004 | 0.58 | 0.92 | 0.29 | 0.63 |
| **SIFT2** | 40.2 ± 35.3 | 39.1 ± 12.9 | 8.9 ± 4.5 | 20.5 ± 5.3 | 0.013 | 0.0009 | 0.61 | 0.94 | 0.64 | **0.79** |

(a) DT-based approach. For each filtering technique the group differences in the mean I_D_ values were detected. However, only for SIFT2 the actual p-value was lower than for the original unfiltered tractogram (0.013 versus 0.014, respectively) - all other techniques showed higher values. The original, non-filtered, tractogram also returned similar results to the filtered ones with regard to the AUC values (0.61, the highest values for filtering were obtained for SIFT2 and LiFE, 0.61 and 0.60, respectively) and to the R^2^ values (0.65 for non-filtered, with SIFT2 and LiFE returning 0.64 and 0.52, respectively, neither being significant). It should be noted, however, that the derived I_D_ values were generally improved, i.e. closer to the known ground truth in comparison to their non-filtered I_D_ counterparts (where estimates of crossing are lower; **Table 2** and **Supplementary Figure A.1**).

(b) For the CSD-based approaches, except for LiFE, all filtered techniques yielded significant group differences for I_D_ and, including LiFE, higher I_D_s for both albinism and controls closer to the ground truth, compared to the unfiltered approaches. The good performance of filtering was also reflected in the highly increased AUC values (with non-filtered tractograms yielding an AUC of 0.75, the COMMIT-SB, COMMIT-SZB and SIFT2 returned AUCs of 0.94, 0.92 and 0.94, respectively). Surprisingly, filtering did not improve the correlation between dMRI and fMRI. Out of 4 tested filtering techniques only SIFT2 returned a significant correlation (R^2^ = 0.79). The best results (highest R^2^ and AUC) were obtained for SIFT2, with COMMIT-SZB being the second best.

### (iv) Robustness of tractography filtering

Given the remarkable similarity of the unfiltered tractograms and the SIFT2-filtered tractograms, it was tested whether applied filtering models are robust to noise in input tractograms. For this purpose, the filtering techniques were applied to SNT (“Streamline Number Targeted”, see Methods) tractograms with forcibly equalized numbers of streamlines as detailed in Methods. This effectively caused for the unfiltered tractograms an overrepresentation of the crossing streamlines, especially in the control population (**Supplementary Table C.1**). The group mean I_D_s were not significantly different, neither for the DT- [median (mean±SEM) I_D_ =36.8% (46.9%±33.3) and 34.7% (28.4%±18.2), for albinism and controls respectively; p=0.093] nor for the CSD-based approaches [median (mean±SEM) decussation index 48.7% (44.2%±7.9) and 48.3% (46.2%±5.4), for albinism and controls respectively; p=0.726]. Subsequent filtering and analysis of filtered SNT tractograms resulted in the following results for the (a) DT- and (b) CSD-based approaches (**Supplementary Table C.1**):

(a) DT-based approach. Already prior to filtering there was a tendency to group differences in the initial mean I_D_ values, which turned significant for all filtered SNT tractograms. With regard to the AUC values, all filtering techniques returned increased AUC values with respect to the unfiltered tractogram. The filtering, however, did not improve the R^2^ values, neither made it the correlation statistically significant.

(b) CSD-based approach. The initial mean I_D_ values were much less distinguishable between groups than for the DT-based approach. This made the group differences after filtering more meaningful. In agreement with the above results (**Suppl. table C.1**), LiFE failed to detect group differences (p=0.58), but this also applied to the SIFT2 results (p=0.68). SIFT2 also returned a lower AUC than without filtering. In contrast, for the COMMIT-SB and the COMMIT-SZB model the group I_D_ differences were significant (0.006 and 0.024, respectively), and the reported AUC values were higher than without filtering (0.83 and 0.78, respectively). All filtering methods failed to detect strong and significant correlations between dMRI and fMRI estimates.

## 4. Discussion

We used diffusion weighted imaging to identify decussation abnormalities at the optic chiasm and compared different tractography methods (DT and CSD) and filtering schemes (LiFE, COMMIT-SZ, COMMIT-SZB, and SIFT2). Even in the case of unfiltered tractograms we reported a significant difference of the decussation index (I_D_) between albinism and controls, which was more pronounced for the CSD than for the DT model. This result is consistent with previous reports that CSD-based tracking can return valuable results that outperform DT-based methods (Franco Pestilli et al. 2014). The ROC analyses for the CSD model suggested its potential as an aid for individualized diagnostics. This was further supported by the better linear correlation of the CSD-derived I_D_ values with the extent of misrouting estimated from fMRI. Further analyses for the tested models and filtering methods consistently confirmed that the CSD model yields better classification accuracies for all investigated approaches. We used two filtering methods with default parameters (LiFE and SIFT2), and a third one that varied in the parameters set to fit the diffusion kernel that optimally matches the dMRI data (COMMIT-SB and -SZB). We replicated previous results demonstrating that the choice of parameters matters (Takemura et al. 2016). We demonstrate that even a single implementation of an evaluation method (COMMIT in our case) can return very different results depending on the choice of parameter set and corresponding quality of fit to the data. Indeed, COMMIT-SB was comparable in performance to the default LiFE and SIFT2. Instead, COMMIT with a kernel model that returned a better fit to the diffusion signal (the -SZB kernel) improved results, with both increased evaluation performance and decreased NRMSE (Supplementary Figure B.1.). These findings are consistent with established knowledge; the better the model fit to the data, the better the evaluation (Franco Pestilli et al. 2014; Takemura et al. 2016; Rokem et al. 2015, 2017).

### Comparison of results for DT with the literature

Our results are in accordance with those reported by Ather et al. (2019) demonstrating that dMRI can detect structural abnormalities of the optic chiasm at a group level. Notably, the findings were reproduced despite several differences in the study design, e.g. acquisition protocols, preprocessing pipeline, implementation of the DT model, tracking algorithms and sample size, which indicates the robustness of the effect. However, the numerical values of mean I_D_ we obtained differ from those previously reported (Ather et al 2019). While Ather et al. (2019) report mean I_D_ for albinism and controls of 42.0% ± 18.7 and 27.8% ± 17.5, respectively, we note that, while our estimates of crossing in albinism show similar values (40.5% ± 40.1), we underestimate crossing in the control group (5.8% ± 4.4). Our values for controls, however, correspond very well with those reported by Roebroeck et al. (Roebroeck et al. 2008). In their study, a ultra-high field (9.4 T) and sub-millimetre resolution (156 × 156 × 312 μm) dMRI analysis of 3 ex-vivo human chiasms using the DT model, they also reported values corresponding to I_D_ ≈ 5%. The discrepancy of the results across studies appears to be linked to different b-values – while Ather et al. used b-value of 1000 s/mm^2^, Roebroeck et al. used b-value of 1584 s/mm^2^, similar to our study (1600 s/mm^2^). Generally speaking, lower b-values preserve more signal originating from extra-axonal compartments, which results in more isotropic tensors. This, in turn, eases tractography and results in the generation of a higher number of streamlines. Those differences would be expected to particularly impact on challenging conditions, such as the mixture of crossing and non-crossing streamlines in the chiasm of controls. Higher b-values, in turn, would severely impair the success ratio of tracking in such conditions, as observed here. More generally, it should be noted that all the estimates of I_D_ inferred from DT analyses of dMRI which were reported thus far [Roebroeck et. al (2008), Ather et. al. (2018)] heavily underestimate the actual ground truth ratio of crossing to non-crossing nerves in the optic chiasm (53:47, respectively) as reported by Kupfer et al. (1967). One of the causes of this is an intrinsic limitation of the DT model, as it assumes only one, dominant direction per voxel and as such is ill-defined for populations of crossing fibers. Consequently, the application of DT in those cases leads to the neglect of valid fibers and erroneous estimates of the primary direction (Wiegell, Larsson, and Wedeen 2000).

### Comparison of results between CSD and DT

Given the established limitations of DT for the crossed fibers as in the chiasm, we extended our study by incorporating a CSD model. This model is believed to be superior to DT in resolving crossing fibers and additionally benefits from the HARDI protocol we used for the acquisition of our data. Accordingly, we found that for non-filtered CSD-based tractograms the mean I_D_ values for albinism and control (41.1 ± 15.2 and 19.2 ± 6.3 %, respectively) were higher and closer to the biological ground truth than those we reported for DT. This is further supported by the cross-modality validation with fMRI-estimated misrouting [R^2^_CSD_ vs R^2^_DT_: 0.83 (p=0.012) and 0.65 (n.s., p=0.052)] and by the ROC-based classification (AUC_CSD_ vs AUC_DT_: 0.75 and 0.61). These findings are in good agreement with theoretical expectations and provide strong support for using models directly incorporating crossed fiber populations in future studies on chiasmal connectivity (Franco Pestilli et al. 2014; J-Donald Tournier et al. 2008; V. J. Wedeen et al. 2008; J-Donald Tournier, Calamante, and Connelly 2012).

### Choice of model parameters and optimal filtering results

We tested only a few filtering methods, not exhaustively as this was not a primary goal of our study. Specifically, two methods were used with default parameters, without optimizing the parameters for the properties of the diffusion data at hand, these methods were LiFE and SIFT2. The COMMIT method was used with two types of parameters (we changed the kernel used for predicting the diffusion, see SZB and SB results). Our results show that the non-optimized methods can return a higher NRMSE in fitting the data, optimized methods instead can fit the data better. As a result of the better fit to the data, filtering improved and allowed higher detection of group differences, i.e. higher AUC-values as compared to the unfiltered tractograms. Even though a full-comparison of all the filtering methods goes beyond the scope of the current work, the results demonstrate that no filtering method is good or bad per-se. Instead careful attention must be taken to assure that the method used in a study achieves a good fit to the diffusion signal, before interpreting and evaluating the results.

### 4.4 Limitations

The study is mainly limited by the challenges posed by the diffusion-weighted imaging of the optic chiasm. Specifically, the small size of the optic chiasm and its location in a highly heterogeneous brain region impact on the quality of dMRI images and subsequent analysis stages, such as tractography. While the design of our acquisition protocol and preprocessing pipeline allowed us to address those challenges, we note that this study aspect may be further improved in the future with emerging methods (Bastiani et al. 2019). In the present work, we provide proof-of-principle results for the potential to identify crossing abnormalities in the chiasm. Our study was limited in the number of participants, and it is possible that future studies with higher participant numbers will allow for refined ROC-analyses and thus suggest diagnostic criteria with higher classification accuracy. For example, in our case, doubling the sample size, would have allowed us to divide the data into a “training” and “test” set to cross-validate the ROC analysis (Franco Pestilli et al. 2014; Rokem et al. 2015; Franco Pestilli 2015). Unfortunately, no datasets of similar quality and scope are currently openly available to perform additional analyses. Such lack of data speaks to the importance of the promotion of open sharing of data and algorithms to advance methods development and scientific understanding (Avesani et al. 2019; Gorgolewski et al. 2017, 2016).

### 4.2 Practical relevance

A key objective of our study was to explore the efficacy of dMRI-based assessment of chiasm integrity and hence its potential as a diagnostic. This is particularly relevant as the identification of chiasmatic abnormalities is a key for the correct diagnosis, especially in phenotypes of albinism with mild pigmentation defects (Montoliu et al. 2014). Functional tests established for this purpose (Apkarian et al. 1983; E. A. H. von D. Hagen, Hoffmann, and Morland 2008) have the disadvantage that they require visual stimulation which in turn relies on the cooperation of the often severely visually impaired participants. While procedures based on purely anatomical MRI procedures do not allow for an individualized identification (Schmitz et al. 2003), our results indicate that dMRI combined with CSD modeling might be able to fill this gap. In fact, as suggested above, testing a greater sample of participants is now required. Although the protocol used in the study is clinically suboptimal due to its length (dMRI and fMRI protocols being, respectively, ∼45 and ∼25 minutes long), the use of simultaneous multi-slice EPI and limiting acquisition of opposing phase-encoding direction to b0 volumes only will shorten dMRI protocol by factor 3-4, i.e., to 10-15 minutes. The full scanning session, including localizer and acquisition of T1-weighted image, would in that case take no longer than an hour.

## 5. Conclusions

We investigated the application of state-of-the-art dMRI to detect optic chiasm abnormalities and report CSD-based models to identify abnormalities with high accuracy (AUC=0.75) and to correlate well with functional (fMRI) measures of the optic nerve misrouting (R^2^=0.83). The classification accuracy (AUC=0.92), as well as veridicality of estimates of crossing strength can be further improved by the application of filtering techniques with optimized parameters (in our case COMMIT-SZB). dMRI combined with CSD-modeling and filtering techniques therefore appear to offer promising approaches for the individualized identification of chiasmatic abnormalities. Moreover, our investigations highlight the great value of the optic chiasm as a test-bed for dMRI methods-optimization. In order to further support these activities, we are in the process of making the data set publicly available for the benefit of the general neuroimaging community.

## Acknowledgements

The authors thank the study participants. This work was supported by European Union’s Horizon 2020 research and innovation programme under the Marie Sklodowska-Curie grant agreement (No. 641805) and by the German research foundation (DFG, HO 2002 10-3) to M.B.H. F.P. was supported by NSF IIS-1636893, NSF BCS-1734853, NSF AOC 1916518, NIH NCATS UL1TR002529, a Microsoft Research Award, Google Cloud Platform, and the Indiana University Areas of Emergent Research initiative “Learning: Brains, Machines, Children.”

---

**Figure A.1.**
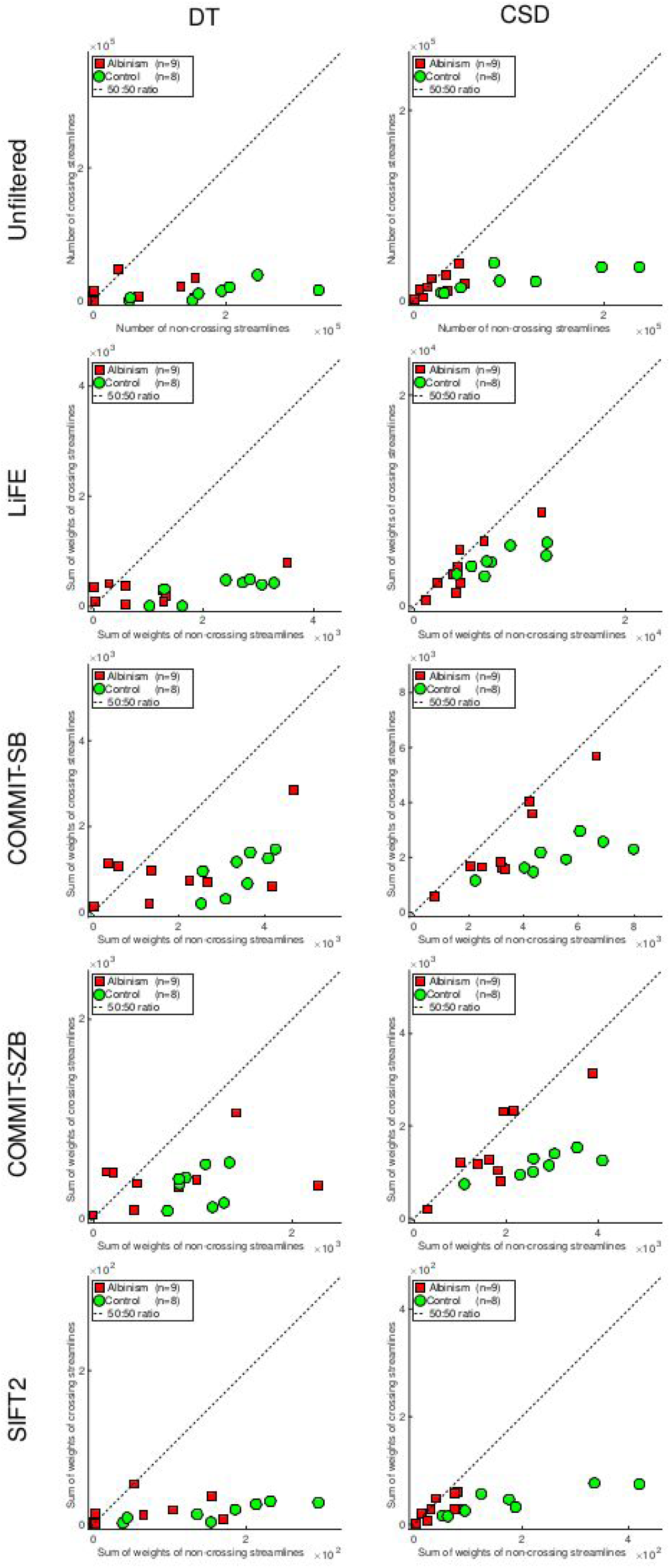
Streamline counts for all tested combinations of diffusion models and filtering methods. Numbers of crossing vs non-crossing streamlines calculated from tractograms based on DT (left column) and CSD (right column) models. Rows correspond to different tested filtering methods as indicated.

**Figure B.1.**
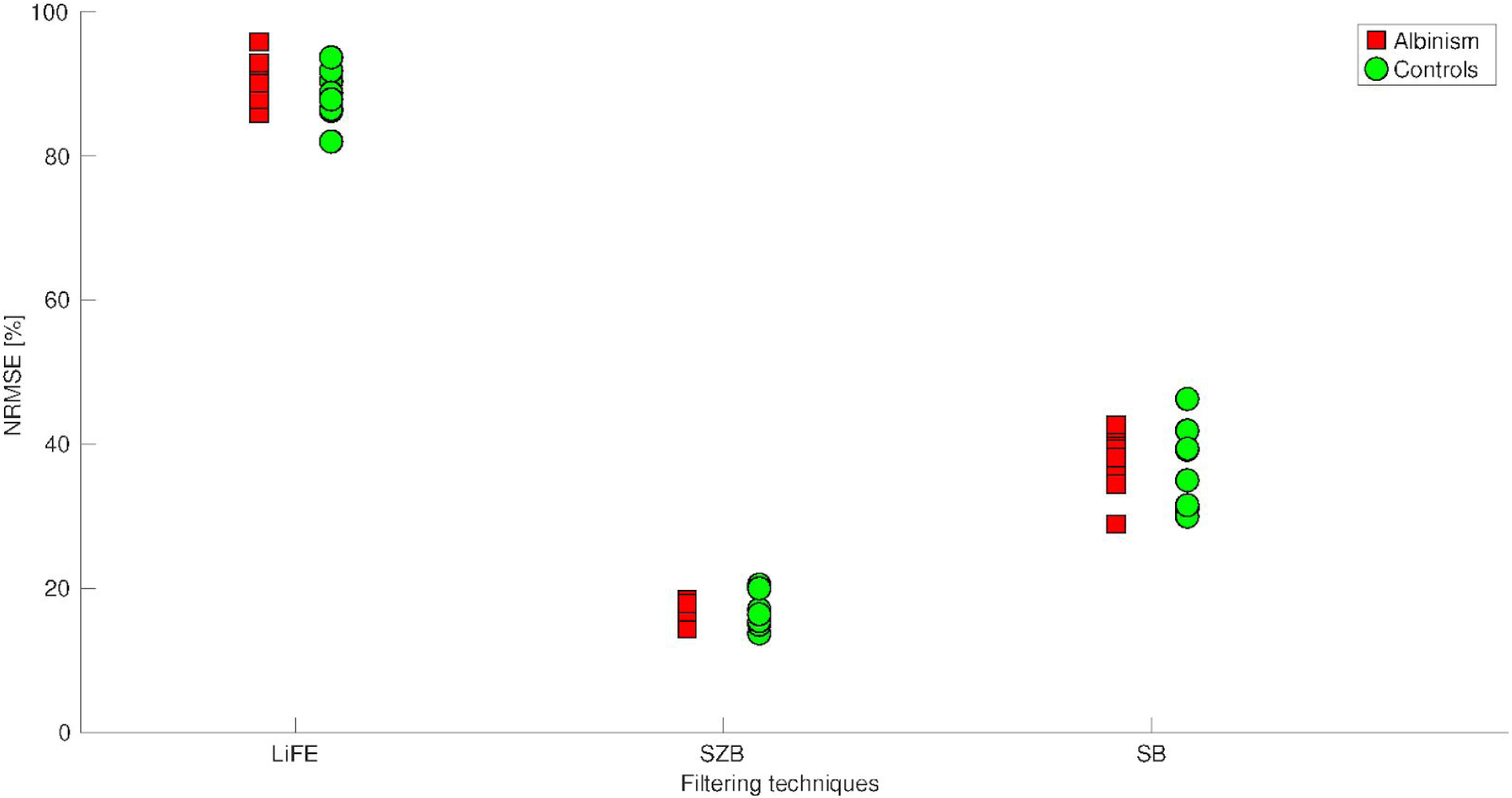
Comparison of NRMSE across tested filtering models for CSD-based tractograms. NRMSE (y-axis) demonstrating quality of the model fit to the data for tested filtering techniques with default parameters of models (x-axis) for both albinism and control group.

**Table C.1.** Results derived from SNT tractograms. Rows describe different filtering techniques, columns describe separate estimates: mean I_D_ calculated for group with albinism, mean I_D_ calculated for controls, p-value derived from statistical testing of equality of mean I_D_s for albinism and controls, AUC calculated from ROC analysis, and R^2^ indicating how much variance in dMRI-based estimates is explained by fMRI results. Each column is divided into two sub-columns, corresponding to DT and CSD signal models.

| | mean $I_D$ for albinism [%] | | mean $I_D$ for controls [%] | | p-value for $I_D$ | | AUC | | $R^2$ | |
| --- | --- | --- | --- | --- | --- | --- | --- | --- | --- | --- |
|  | DT | CSD | DT | CSD | DT | CSD | DT | CSD | DT | CSD |
| **Unfiltered** | 46.9 ± 33.3 | 44.2 ± 7.9 | 28.4 ± 18.2 | 46.2 ± 5.4 | 0.09 | 0.72 | 0.51 | 0.54 | 0.53 | 0.09 |
| **LiFE** | 43.3 ± 30.5 | 46.3 ± 10.8 | 21.5 ± 7.3 | 47.1 ± 4.7 | 0.03 | 0.58 | 0.60 | 0.63 | 0.29 | 0.11 |
| **COMMIT - SB** | 44.7 ± 25.5 | 45.4 ± 5.8 | 24.2 ± 8.4 | 38.8 ± 3.1 | 0.02 | 0.006 | 0.60 | 0.83 | 0.25 | 0.0003 |
| **COMMIT- SZB** | 49.8 ± 23.2 | 49.0 ± 8.5 | 25.9 ± 12.3 | 42.0 ± 3.5 | 0.01 | 0.024 | 0.64 | 0.78 | 0.31 | 0.18 |
| **SIFT2** | 45.2 ± 29.7 | 41.8 ± 8.4 | 24.5 ± 12.4 | 43.4 ± 5.0 | 0.04 | 0.68 | 0.56 | 0.50 | 0.46 | 0.06 |

