## Supplementary figures and images for "Quantifying nerve decussation abnormalities in the optic chiasm"

### Supplementary Figure A.1

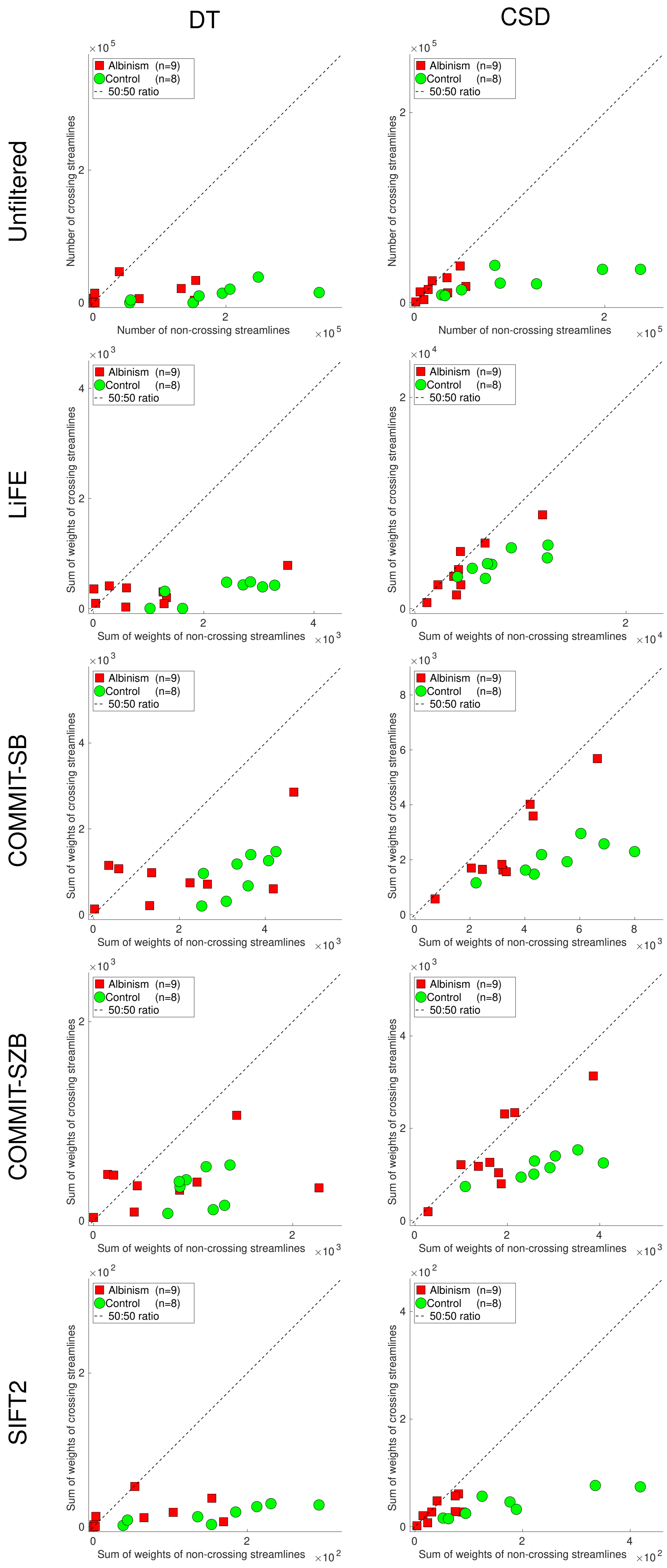

### Supplementary Figure B.1

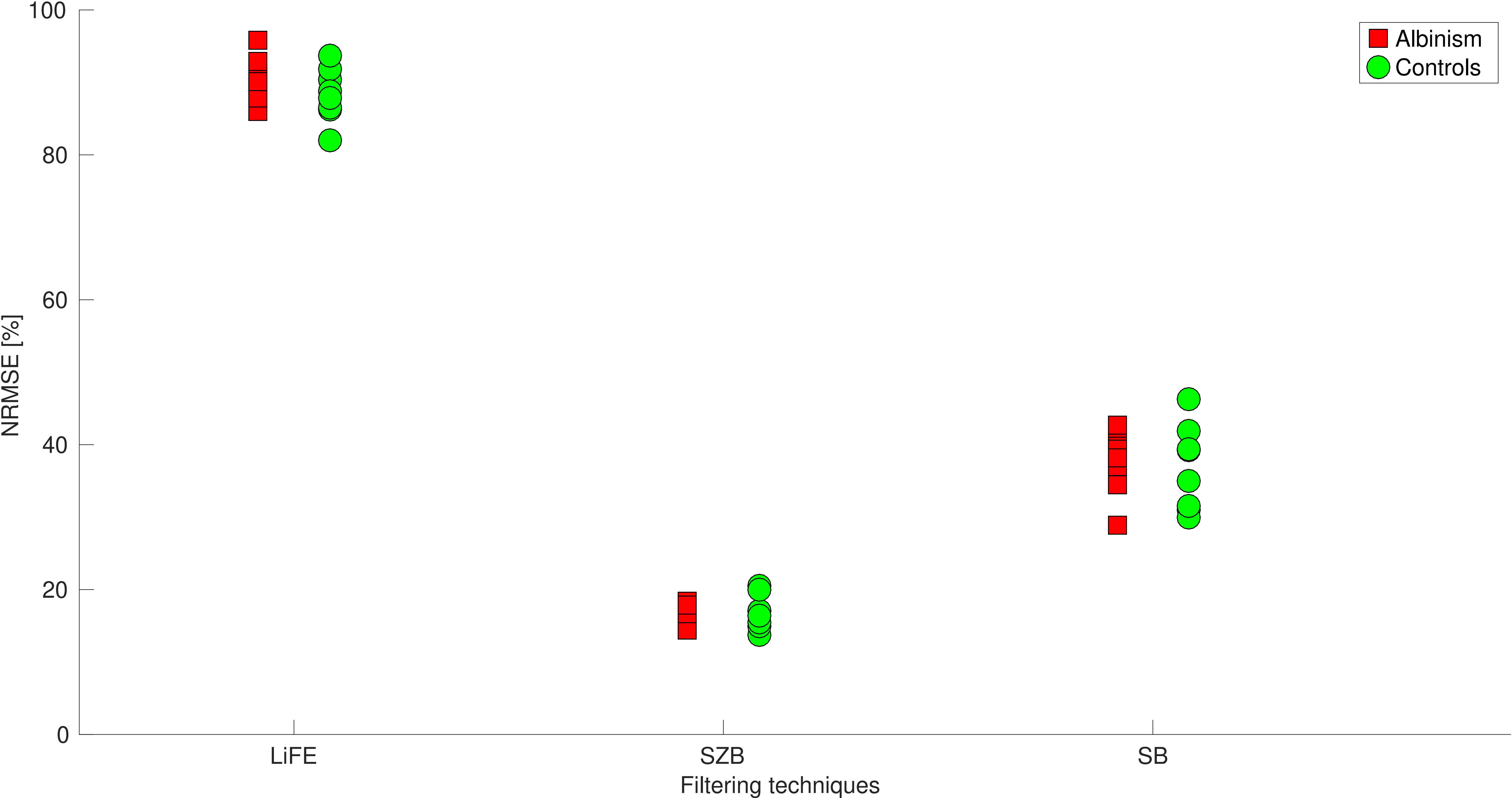
